# Trait-based predictors of genomic diversity within freshwater fish communities of the Ozarks

**DOI:** 10.1101/2022.10.26.513843

**Authors:** Zachery D. Zbinden, Marlis R. Douglas, Tyler K. Chafin, Michael E. Douglas

## Abstract

Phenotypic traits modulate the fate of species interactions with one another and the environment; thus, traits directly shape the past, present, and future evolutionary trajectories of populations. As such, distinct species-specific responses to a shifting environment are widely documented in the form of distinct genetic signatures, i.e., genetic diversity, reflecting differential responses over time. While the link between genetic diversity and phenotypic traits is seemingly fundamental, it has been challenging to establish unequivocally. Across an exemplar freshwater fish metacommunity, we employ phenotypic traits to test if they are significantly related to observed genetic patterns among species. Associated traits were then used to construct trait-based predictive models of genetic diversity. We collected representative constituents of a freshwater fish community (*N*=31 species) sampled across 75 sites within the White River Basin (Ozark Plateau, USA). For each species, we derived three genetic diversity indices (=*H*_S_/*H*_T_/*G*”_ST_) from SNP data (*N*=2,000 loci) and assessed 28 phenotypic traits related to morphology, life history, and ecology. We identified a series of traits (*N*=2–5, depending upon the index) strongly associated with facets of genetic diversity. These were subsequently applied in predictive models that explained 31–68% of the genetic variability across species, suggesting a potential utility as an imputation tactic for data-deficient species. Our approach effectively linked species-specific traits with genetic diversity within and among populations, thereby further clarifying correlations between contemporary ecological processes, as modulated by species traits, and long-term evolutionary trajectories.

## 1 INTRODUCTION

How species interact with their abiotic and biotic environment ultimately determines the evolutionary trajectory of populations (Hand et al., 2015; Hutchinson, 1965). Linking ecology and evolution remains a central challenge in biology (Avise et al., 2016). One way to link them is through the relationship between species’ phenotypic traits and genetic diversity (Bohonak, 1999). Phenotypic characteristics mediate how species interact with abiotic and biotic environmental factors, which in turn influence population size and connectivity fluctuations through time. Therefore, traits also play a role in the evolution of populations and should show strong relationships with different evolutionary metrics (i.e., genetic diversity; Meirmans et al., 2011). Here, ‘genetic diversity’ can be further partitioned into three distinct ‘facets,’ i.e., within-site diversity (α), among-site diversity (β), and total diversity (γ) (Sherwin et al., 2017). Parsing the relationship between traits and genetic diversity to uncover generalities requires a comparative approach (López-Uribe et al., 2019; Selkoe et al., 2014).

We posit that the most insight to be gained is at the community level, e.g., via landscape community genetics (Hand et al., 2015). By community-level, we mean simultaneously evaluating genetic population structure within multiple, co-distributed species (Leibold et al., 2004; Rissler, 2016). At this level, the study of ecology and evolution is enhanced by quantifiable variation among species that can be leveraged to dissect the relationship between species-level characteristics and their emergent qualities. Applying this manner of trait-based approach within an explicitly comparative framework acts as a ‘window’ through which to view the associative link between traits shaping ecology and evolutionary history, for example, as demonstrated by statistical associations between genetic diversity and the traits in question.

Theory provides predictions that might be tested by applying such a ‘trait-based’ approach within a comparative framework. In an ecological sense, life history influences demography and population size, while dispersal-related traits tied to morphology and ecology initiate spatial connectivity. These processes, accumulated over time, impact genetic diversity through various effects on evolutionary processes - smaller populations rapidly lose genetic variation to drift (Lynch & Lande, 1998), while those with limited connectivity diverge more rapidly in a genetic sense (Wright, 1931). Thus, the link between traits and genetic diversity is substantial and has been empirically demonstrated in animals (Bohonak, 1999), plants (Duminil et al., 2007; Loveless & Hamrick, 1984; Meirmans et al., 2011), marine organisms (Kelly & Palumbi, 2010; Riginos et al., 2014), and birds (Burney & Brumfield, 2009). These previous studies tested the relationships between traits and genetic diversity and, in so doing, supported the link between ecology and evolution. However, trait diagnosis and predictability remain inconsistent (Riginos et al., 2014), a conclusion not surprisingly derived from hundreds of different population-level studies across many different biogeographical regions (but see: Meirmans et al., 2011; Selkoe et al., 2014; Sousa-Santos et al., 2016). The search for targeted associations becomes arduous and muddled as numerous confounding factors are incorporated (Kelly & Palumbi, 2010). Additionally, meta-analyses are often forced to incorporate legacy markers with lower information content, e.g., isozymes, AFLPs, or microsatellites. A community-level approach that genotypes individuals and estimates genomic diversity using next-generation sequencing technology might allow for a potentially more robust link between traits and diversity and reduce inconsistencies among projected studies.

If life history and dispersal-related traits are causally associated with genetic diversity, then diversity could be predicted in other species by modeling their traits; this could be valuable because genetic diversity represents the variation available for evolution to act upon and serves as an indicator of population persistence (Jump et al., 2009). The spatial structure of genetic diversity further imposes constraints on a species’ ability to adapt to a changing environment (López-Uribe et al., 2019). Many efforts have aimed to assess the genetic diversity of threatened species, although over 70% have yet to be assessed (Bachman et al., 2019; Hogg et al., 2022), underscoring the value of a predictive framework. Understanding levels of standing genetic variation and its structure is critical for managing declining species (Willoughby et al., 2015).

However, suppose we could forecast which species will become threatened in the future based on genetic diversity indicators – as a proxy for population persistence. We could then take proactive measures to bolster such species against decline (Lunney et al., 2004; Martinez, Willoughby, & Christie, 2018). These predictions can guide efforts to prioritize species of conservation concern and focus targeted efforts aimed at further data collection. Rather than conducting population genetic studies on hundreds or thousands of species, which would be costly and require technical skill and necessary infrastructure, we could make accurate predictions of critical conservation indicators by assessing a few dozen species representative of a regional metacommunity and its trait variation.

The primary focus of this study was to test the hypothesis that species-specific traits within a freshwater fish community are related to genetic diversity. We hypothesized that essential traits would be related to morphology, life history, and ecology, as these are most likely to impact population size, reproduction, and dispersal. Little work has been done to assess the relationship between traits and genetic diversity for freshwater fishes (but see Martinez et al., 2018; Mitton & Lewis, 1989; Pilger et al., 2017; Sousa-Santos et al., 2016). Secondly, we were interested in which traits are most strongly associated with diversity and what inferences, i.e., mechanisms promoting genetic diversity, could be established from these relationships. Finally, we constructed predictive models of different genetic diversity indices based on parsimonious sets of traits.

## 2 METHODS

### 2.1 The study region

The focal region for our study is the Ozark Plateau (Figure 1), specifically the White River Basin, which drains 71,911 km^2^ as a tributary to the Mississippi River. Like the Appalachian Highlands, the Ozark Highlands served as a glacial refugium during the Pleistocene (Mayden, 1988). Its long temporal span of geologic stasis has acted to equilibrate gene flow and genetic drift, thus serving as a ‘natural laboratory’ for biodiversity diversification (Hutchison & Templeton, 1999). Here, we note an important contrast with more northern latitudes, where cyclical bottlenecks and subsequent expansions (e.g., as a by-product of glacial/ interglacial cycles) have effectively recast population genetic signatures of resident taxa (Cammen et al., 2018). This presents a problem in that genetic signatures may be effectively ‘masked’ by non-equilibrium demographic processes driven by geomorphic disturbances. Thus, we expect such patterns to be more accurately preserved within glacial refugia (i.e., including the Ozark Highlands), thus more amenable to direct interrogation within a hypothesis-testing framework, as herein.

**FIGURE 1.**
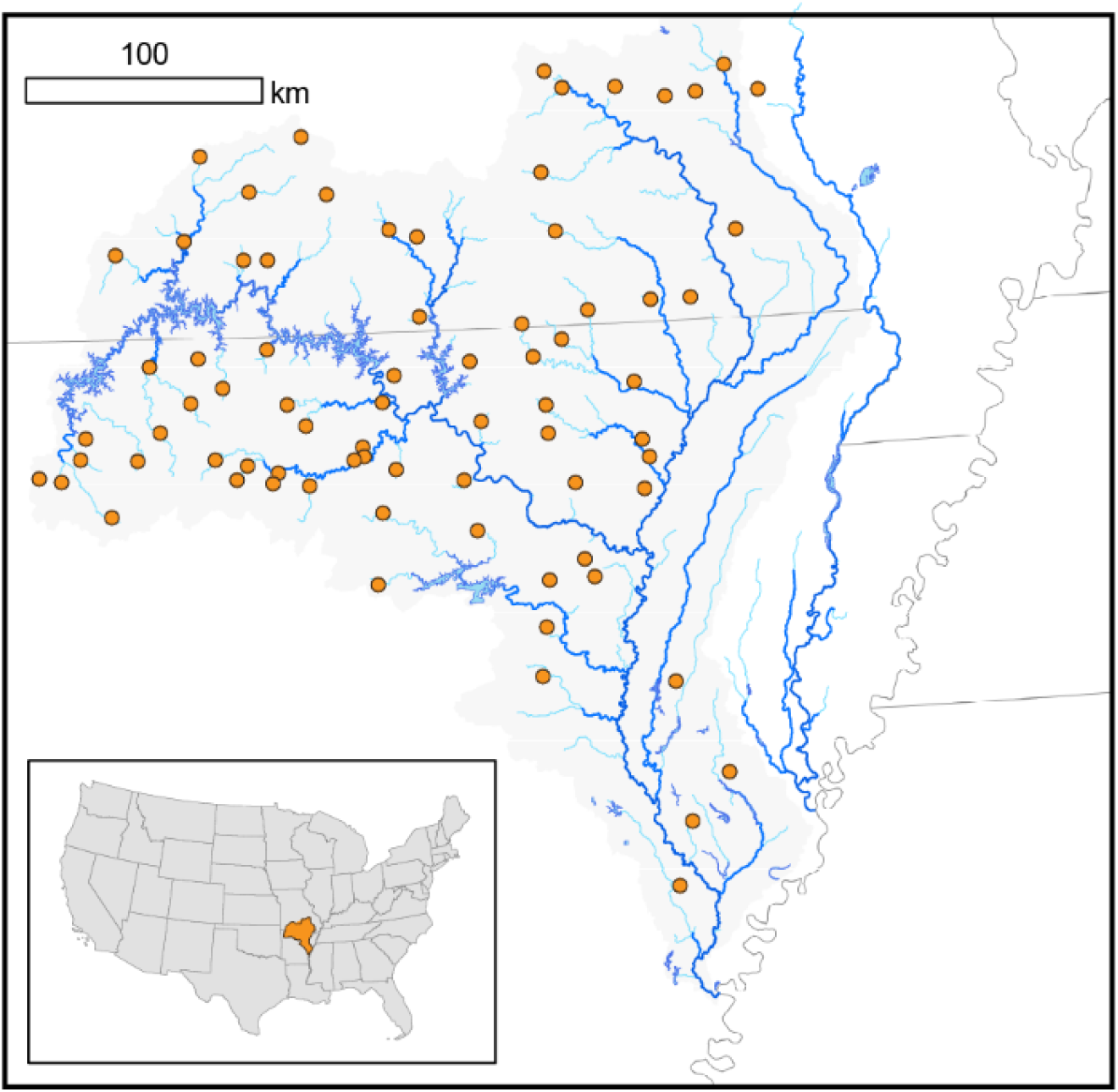
Freshwater fish species (N=31; Table 1) were collected at *N*=75 sampling locations primarily across the White River Basin (Ozark Plateau, USA). Individuals were genotyped at N = 2000 SNPs to quantify facets of genetic diversity.

### 2.2 Sampling

Fishes were collected in wadable streams using seines during low flows between June 2017 and September 2018. Sampling was approved by the University of Arkansas Institutional Animal Care and Use Committee (IACUC: #17077), with collecting permits from Arkansas Game & Fish Commission (#020120191), Missouri Department of Wildlife Conservation (#18136), and U.S. National Parks Service (NPS: Buffalo River Permit; BUFF-2017-SCI-0013). Fish were euthanized by immersion in tricaine methanesulfonate (MS-222) at a concentration of 500 mg/L, buffered to pH=7, and then preserved in 95% ethanol. Species diagnosis occurred in the laboratory, as augmented by Eschmeyer’s Catalog of Fishes (Fricke et al., 2022). The right pectoral fin was removed and stored in 95% ethanol at −20 °C prior to subsequent DNA extraction using QIAamp Fast DNA Tissue kits (Qiagen, Inc.) following the manufacturer’s protocols.

### 2.2 Genomic data collection and filtering

To estimate genetic diversity indices, we developed genotypic alignments for each species using a double-digest restriction site-associated DNA (ddRAD) sequencing procedure (Peterson et al., 2012). Our standard procedures were appropriately modified and previously reported in detail (Zbinden et al., 2022a, Zbinden et al., 2022b).

Genomic DNA was first isolated and then digested with high-fidelity restriction enzymes *Msp*I and *Pst*I. Bead-purified samples were quantified by fluorometry (Qubit; Thermo Fisher Scientific), standardized to 100 ng DNA, and then ligated with custom adapters containing in-line identifying barcodes. Samples were pooled into sets of 48 and subsequently size-selected from 326–426 bp (including adapter length). Illumina adapters with unique i7 indices were added via 12-cycle PCR with Phusion high-fidelity DNA polymerase (New England Biolabs, Inc.). Three libraries were pooled per lane and sequenced single-end on an Illumina HiSeq 4000 platform (100bp read length; Genomics & Cell Characterization Core Facility; University of Oregon, Eugene).

Raw Illumina reads were demultiplexed, clustered, filtered, and aligned with IPYRAD v.0.9.62 (Eaton & Overcast, 2020). Demultiplexing criteria allowed no more than a single barcode mismatch and individuals with extremely low reads 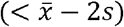 were removed. Next, individuals were screened for misidentifications and putative hybrids (see Zbinden et al., 2022a), the former re-assigned and the latter removed. Raw sequence reads were partitioned by species and aligned *de novo*. Clusters of homologous loci were assembled using an 85% identity threshold. Putative homologs were removed according to the following criteria: <20x and >500x coverage per individual; >5% of consensus nucleotides ambiguous; >20% of nucleotides polymorphic; >8 indels present; or presence in <15% of individuals. Paralogs were identified (and removed) as those clusters with either >2 alleles per site in consensus sequence or excessive heterozygosity (>5% of consensus bases or >50% heterozygosity/site among individuals).

Biallelic SNP panels were then visualized for each species, with additional filtering implemented with the R package RADIATOR (Gosselin, 2020). To ensure high data quality, loci were removed if: Monomorphic; minor allele frequency <3%; Mean coverage <20 or >200; Missing data >30%; SNP position on read >91; and significant HWE deviation in one or more sampling sites (*α* = 0.0001). Linkage disequilibrium was reduced by retaining but one SNP per locus (i.e., that which maximized minor allele count). Finally, singleton individuals/species/sampling site were removed, as were those with >75% missing data in the filtered panel. We standardized panels to eliminate bias caused by differences in the number of loci between species by randomly sampling 2000 SNPs per species panel. However, genetic diversity indices were almost perfectly correlated (*r* > 0.99) whether generated from full *versus* standardized panels.

### 2.3 Genetic diversity

We calculated three facets of genetic diversity for each species: Within-site diversity (α), among-site diversity (β), and total diversity within and among sites (γ) (Sherwin et al., 2017). Three indices were employed to represent these facets: *H*_S_, the average gene diversity (heterozygosity) within sites (Nei, 1973; Nei & Chesser, 1983); *H*_T_, total gene diversity across all sites (Nei, 1973; Nei & Chesser, 1983); and *G*”_ST_, the unbiased fixation index (Meirmans & Hedrick, 2011), which captures the relationship between the two. We chose *G*”_ST_ because it is a more appropriate metric for comparison among species, and it does not underestimate differences when the number of sampled populations is small (Meirmans & Hedrick, 2011). These genetic diversity indices were calculated using R Statistical Software (R Core Team, 2022) via SNP panels formatted as ‘genind’ objects (Jombart, 2008), with calculations performed by the R package MMOD (Winter, 2012). Per Nei (1973), *H*_S_ and *H*_T_ were first averaged across loci before computing *G*”_ST_ as 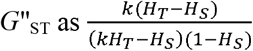 (eq. 4 from Meirmans & Hedrick, 2011, as implemented MMOD).

### 2.4 Explanatory factors

#### 2.4.1 Covariates

Although our goal was to relate species’ phenotypic traits with genetic diversity, we surmised that differences in sampling among species, e.g., number of sites, could also increase variability. Therefore, we included four covariates for each species in our exploratory analysis, to include: the number of individuals in the analysis (=Nindiv), the number of sites at which those individuals occurred (=Nsites), the mean number of individuals analyzed per occupied site (=Mean_Ind_Site), and the median value of pairwise river distances between all occupied sites (=Median_RivDist).

#### 2.4.2 Phylogeny

Closely related species tend to be more similar, which can violate assumptions of independence in our models, i.e., phylogenetic autocorrelation (Felsenstein, 1985). However, removing autocorrelation entirely from analyses can eliminate signals of potential interest (Legendre, 1993; Peres-Neto, 2006). Therefore, we explored phylogenetic autocorrelation using variation partitioning (Borcard et al., 1992), which allowed us to test the effects of traits with and without phylogenetic autocorrelation and determine the potential role of phylogenetically correlated traits.

A phylogeny of the study species was obtained using a previously constructed tree of ray-finned fishes based on a 27-gene multi-locus alignment (Rabosky et al., 2018). Our backbone tree was limited to the species collected in this study (via the R package `fishtree`; Chang et al., 2019). We decomposed the topology using phylogenetic eigenvector regression (PVR; Diniz-Filho et al., 1998) to create a matrix of *N*-1 eigenvectors for a given phylogeny (where *N*=number of species). Principal coordinate analysis decomposed phylogenetic distances among tips into vectors representing the phylogenetic topology, with the first (e.g., EV1) representing deeper splits and more recent splits by the latter (e.g., EV20) (Diniz-Filho et al., 2012). We tested the relationships between these eigenvectors and genetic diversity indices using the approaches described below.

#### 2.4.3 Traits

We assembled a data set of 28 species-level traits broadly related to morphology, life history, and ecology. Phenotypic characteristics within each of these categories can leave an indelible imprint on genetic diversity by impacting dispersal and population demography and, in turn, gene flow and genetic drift. Traits were gathered from three primary databases: FishTraits, a public database for North American freshwater fishes (Frimpong & Angermeier, 2009); FISHMORPH, a global database on morphological traits of freshwater fishes; and an unpublished database (J.D. Olden, unpublished data, 2021) used in previous publications (Giam & Olden, 2016; Mims et al., 2010). Robison & Buchanan (2020) was consulted to corroborate/adjudicate disagreements [Supplementary Material (S1) and online data repository (https://osf.io/837vj/)].

### 2.5 Analyses

#### 2.5.1 Data reduction

We developed three response variables (i.e., indices of genetic diversity) that we hypothesize as related to our predictor variables: (a) Covariates; (b) phylogenetic eigenvectors; and (c) species-level traits. We analyzed each response variable separately using an identical framework (Figure 2) as follows: We first applied vector fitting (R package `vegan`; Oksanen et al., 2020) to test each predictor set against diversity indices. Those variables exhibiting a significant relationship (α < 0.05) were retained and used in separate stepwise forward-selection procedures (i.e., for each diversity index) to yield parsimonious models based on Akaike Information Criteria (AIC) (Oksanen et al., 2020). If covariates explained significant variation in a genetic index (only the case for *H*_T_), those effects were removed using residuals extracted from a generalized linear model (GLM) between the genetic index and significant covariates. This overall approach, along with the multiple regression used below, has yielded reasonable estimates of influential predictor variables via simulation and is robust to the collinearity of predictors (Smith et al., 2009).

**FIGURE 2.**
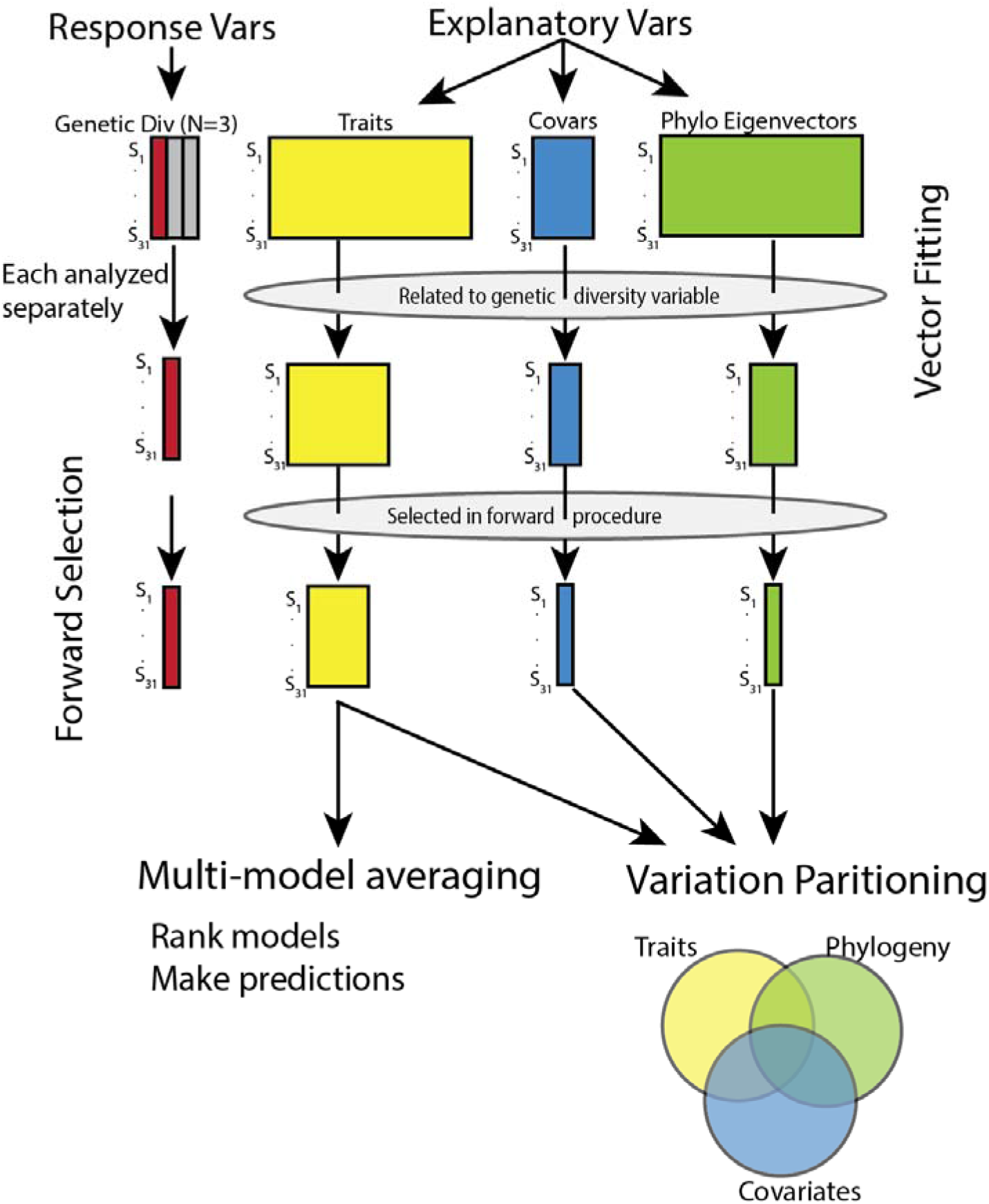
Overview of the framework for hypothesis testing and the model building used in the evaluation of freshwater fish species (N=31; Table 1) collected at *N*=75 sampling locations primarily across the White River Basin (Ozark Plateau, USA).

#### 2.5.2 Variance partitioning

We estimated the variation in genetic diversity explained by each predictor set after accounting for variation explained by the other sets. We partitioned the variation among species across reduced sets of predictor variables for each genetic diversity index to test for significant relationships. Partial multiple regressions (variation partitioning; Borcard et al., 1992; Legendre & Legendre, 2012; Peres-Neto et al., 2006) allowed us to elucidate the variation of each genetic index explained by traits, phylogeny, and phylogenetically correlated traits. Covariates were not included in that they were either not significantly associated with genetic diversity (*H*_S_, *G*”_ST_), or their significant effects had previously been removed per above (*H*_T_). Results were presented as Venn diagrams depicting the adjusted *R*^2^, i.e., the extent of variation explained by predictors. We tested our fractions of variation, e.g., “pure trait” variation after adjustment for phylogeny, using *N=9,999* permutations (Anderson & Legendre, 1999).

#### 2.5.3 Multi-model averaging

Reduced trait sets were used as ‘global’ models to generate all possible combinations, i.e., subsets, with the different models ranked according to second-order Akaike Information Criteria (AIC_C_) (R package MuMIn; Barton, 2022). We derived coefficients for each trait based on model averaging via information criteria to produce predictions based on trait inputs. As a demonstration, we used the models to predict genetic diversity for Slender Madtom (*Noturus exilis* Nelson, 1876), a small-bodied benthic-dwelling catfish in the White River Basin (as an aside, it was not sampled extensively enough to be included in formal analyses; i.e., ≥5sites with ≥2 individuals). We used trait values gathered from the abovementioned sources and model coefficients to predict its genetic diversity values. Slender Madtom SNP genotypes were used to estimate genetic diversity indices (generated in the same manner as for the other species and standardized to *N*=2000 SNPs). These estimates allowed us to compare and contrast predicted *versus* observed genetic diversity indices, thus acting as a ‘proof-of-concept’ regarding their predictive utility when applied to data-deficient species.

## 3 RESULTS

We collected freshwater fish from *N*=75 locations (Figure 1) and analyzed *N*=2,861 individuals representing 31 fish species, as genotyped across standardized SNP panels generated with ddRAD. Each panel was based on 15–358 individuals collected across 5–50 sampling sites (Table 1). The mean number of individuals/species/site =5.1. Mean within-site gene diversity (=*H*_S_) ranged from 0.024–0.283 (x□=0.159; *s*=0.056); total gene diversity (=*H*_T_) ranged from 0.138–0.350 (x□=0.221; *s*=0.053); and among-site diversity (=*G*”_ST_) spanned from 0.022–0.965 (*x*□=0.303; *s*=0.253) [Supplementary Material (S2–S3)].

**TABLE 1.**
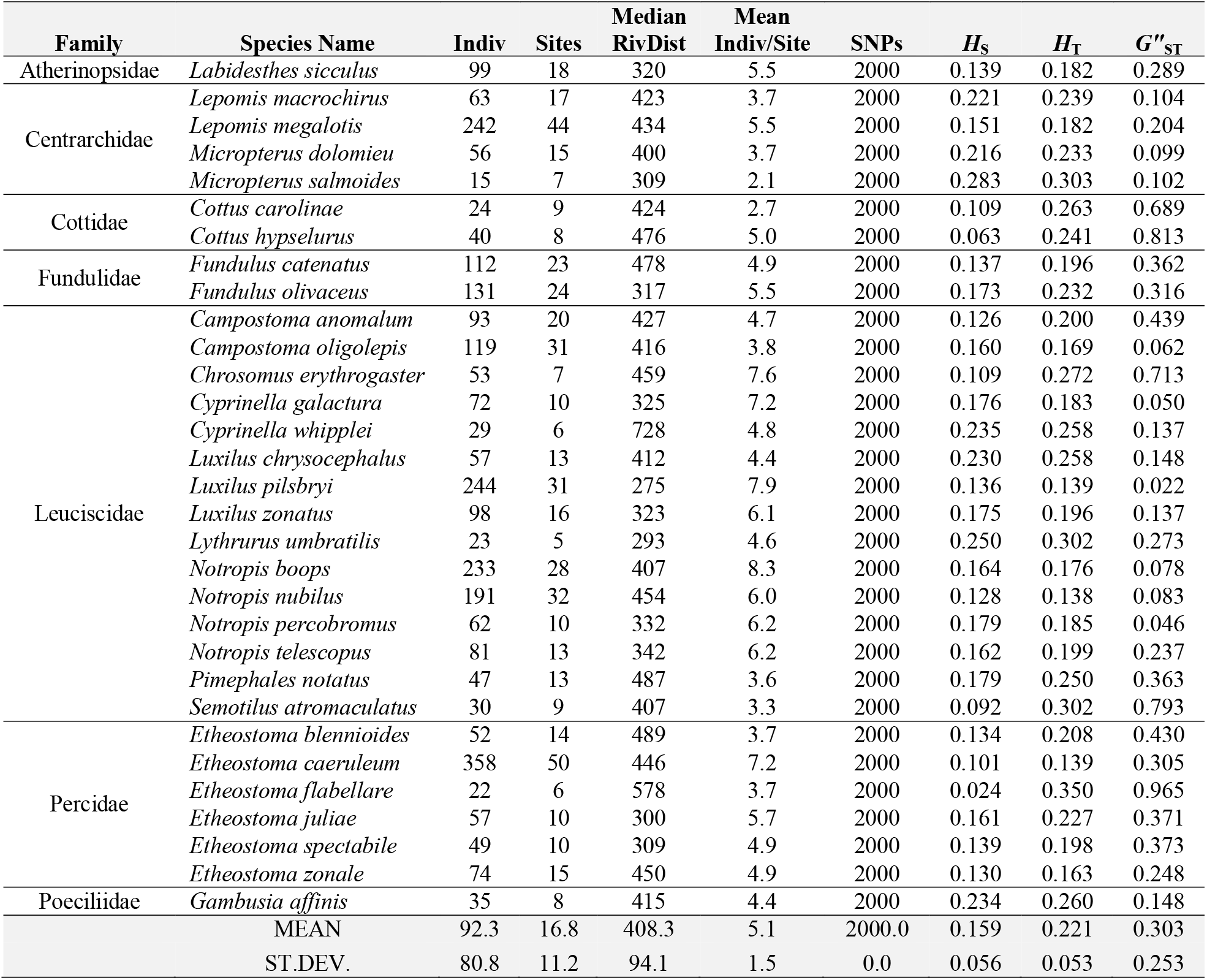
Freshwater fish species (N=31; listed by Family and Species) were collected across the White River Basin (Ozark Plateau, USA). SNP panels resulted from genotyping individuals using ddRAD seq. Panels of randomly selected SNPs (N=2000) were used to calculate average site genetic diversity (*H*_S_), total genetic diversity (*H*_T_), and global genetic fixation/differentiation index (*G*”_ST_). Four covariates are shown: Indiv = Total number of individuals/species analyzed; Sites = Total number of sampling locations/species analyzed; MedianRivDist = Median of pairwise river distance between analyzed set of sites; Mean Indiv/Site = Mean number of individuals analyzed across sites where a species occurred.

We gathered 28 traits related to morphology, life history, and ecology [Table 2; Supplementary Material (S1 & S4)]. We examined relationships between genetic diversity and four covariates (N individuals, N sites, individuals/site, and median river distance between sites). Total gene diversity (*H*_T_) was significantly related to N individuals, N sites, and Mean individuals/site, i.e., sample size (Table 3). These three covariates explained ~62% of the variation in *H*_T_, and forward-selection revealed that N individuals alone could explain ~50%. This finding is not surprising, given that *H_T_* is a sample size-corrected measure (Nei & Chesser, 1983). As such, estimated *H*_T_ is expected to decrease with sample size. The remaining results related to total gene diversity, *H*_T_, are therefore based on the residuals of a linear model (*H*_T_ ~ N individuals) to remove the effect of sampling variability. Although both within-site diversity (*H*_S_) and among-site diversity (*G*”_ST_) have direct mathematical relationships with facets of sample size (Meirmans & Hedrick, 2003; Nei & Chesser, 1983), neither showed any statistical relationship here, thus were analyzed directly in downstream analyses (rather than as residuals).

**TABLE 2.**
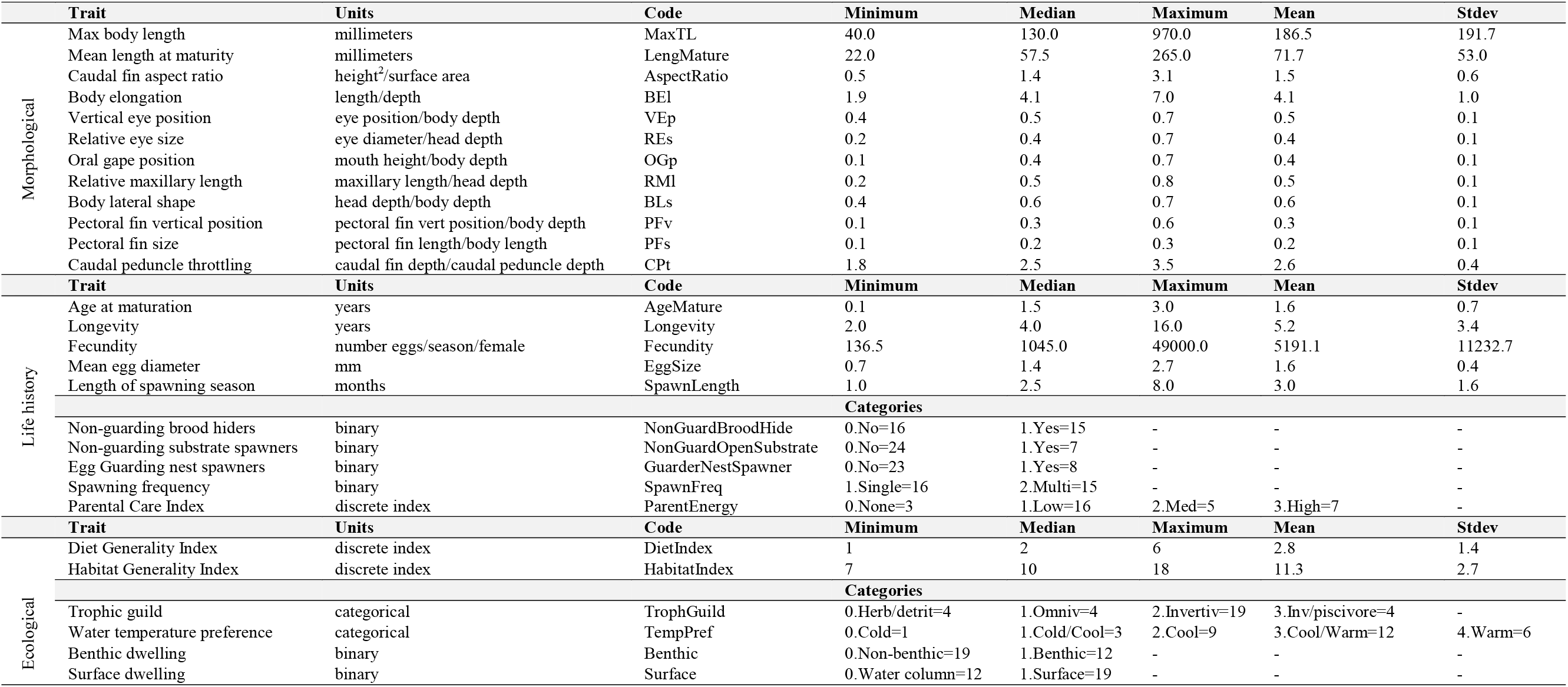
Summary statistics for trait data compiled for freshwater fish species (*N*=31; Table 1) collected across the White River Basin (Ozark Plateau, USA). Traits are grouped as Morphological, Life history, and Ecological and are hypothesized to relate to variation in facets of genetic diversity among species. Trait = Trait definition; Units = As measured; Code = Trait acronym; Categories = 0 (not present)/ 1 (present); Stdev = Standard deviation.

**TABLE 3.** Summary of analyses testing the relationship between traits and three facets of genetic diversity quantified using SNPs (N=2000) genotyped for freshwater fish species (N=31; Table 1) collected across the White River Basin (Ozark Plateau, USA). Genetic diversity facets quantified for all species included: Within-site diversity (*H*_S_), total diversity (*H*_T_), and among-site diversity (*G*”_ST_). We tested relationships between genetic diversity variation among species and: covariates, traits (Table 2), and phylogenetic relatedness. Results for those tests are provided here. For each facet (*H*_S_/*H*_T_/*G*”_ST_) and each predictor variable set, variables were tested for a relationship with a given facet using vector fitting (significant variables denoted “Fitted Vars”). These were then subjected to a forward-selection procedure (“Selected Vars”). The adjusted coefficient of determination for the selected variables (“Selected *R*^2^”) is based on multiple regression, and all values shown are significant (α < 0.05).

| | | $H_S$ | $H_T$ | $G''_{ST}$ |
| --- | --- | --- | --- | --- |
|  | Fitted Vars | none | N indiv; N sites; Mean Indiv/Site | none |
| Covariates | Selected Vars | - | Nindiv | - |
| | Selected $R^2$ | - | 0.5 | - |
| Traits | Fitted Vars | MaxTL; OGp; Fecundity; EggSize; SpawnLength;<br>TempPref; Benthic; Surface | BEI; BLs; ParentEnergy;<br>GuarderNestSpawner | OGp; PFv; CPt; EggSize; Benthic;<br>Surface |
|  | Selected Vars | Surface; TempPref; EggSize; MaxTL; SpawnLength | BLs; GuarderNestSpawner | Surface; EggSize |
| | Selected $R^2$ | 0.68 | 0.31 | 0.31 |
| Phylogeny | Fitted Vars | EV2 | EV18 | EV4; EV6 |
|  | Selected Vars | EV2 | EV18 | EV4; EV6 |
| | Selected $R^2$ | 0.15 | 0.36 | 0.36 |

Each facet of genetic diversity was significantly related to some phenotypic traits: *H*_S_=8 traits; *H*_T_=4; *G*”_ST_=6 (Table 3). These reduced sets explained 26–66% of the variation within diversity (Table 3). Following forward-selection, a parsimonious trait model was derived for each index and involved fewer traits: *H*_S_=5 traits; *H*_T_=2; *G*”_ST_=2 (Table 3). These explained similar amounts of variation (31–68%), as did the vector-fitted sets (Table 3).

Each facet of genetic diversity was also significantly related to phylogenetic distance (Table 3). Only one or two phylogenetic eigenvectors were significantly related to each genetic diversity index but explained between 15–36% of the variance. In each case, those phylogenetic eigenvectors that were significant in the vector fitting reduction step were also selected in forward-selection (Figure 2; Table 3). Relationships between each genetic diversity index and its corresponding vector-fitted variables are found in Supplementary Material (S5–S7).

After accounting for phylogenetic autocorrelation, significant variance in genetic diversity was explained by partial multiple regression among traits (Figure 3). Total variation (traits and phylogeny) ranged from 43–68%, whereas variance explained “purely” by traits ranged from 6–52%. Phylogeny explained from 0–18% of the variation. Finally, phylogenetically correlated traits explained 16–25%.

**FIGURE 3.**
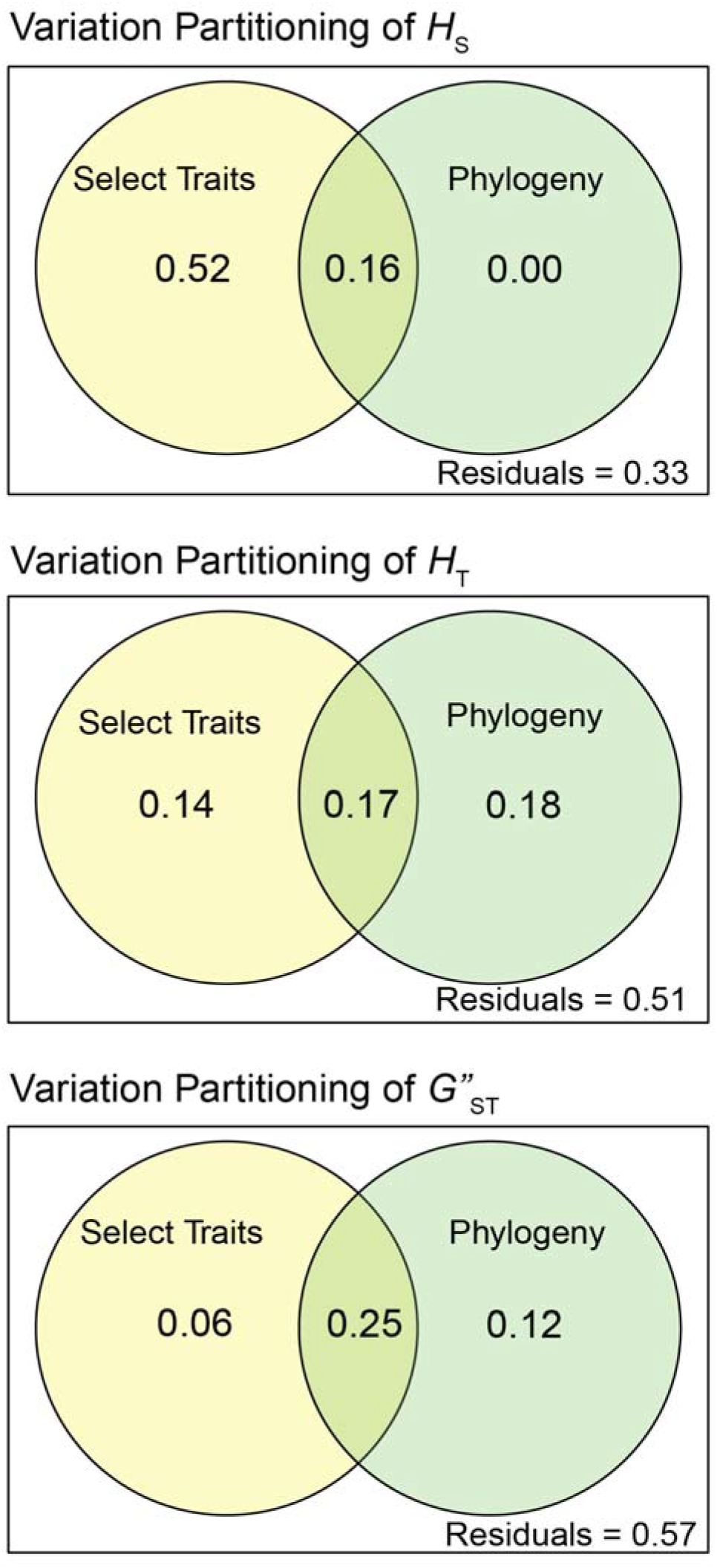
Variation across freshwater fish species (N=31; Table 1) collected at *N*=75 sampling locations primarily across the White River Basin (Ozark Plateau, USA) for each of three genetic diversity indices: Within-site diversity (*H*_S_); total diversity (*H*_T_); and among-site diversity (*G*”_ST_). Each index was calculated by species based on single nucleotide polymorphisms (N=2000). Interspecific variability within each index was partitioned among selected sets of traits (Table 2) and phylogenetic eigenvectors. Values represent adjusted *R*^2^ or the percentage of variation explained by either [A] select traits, [C] phylogeny, [B] both/phylogenetically correlated traits, or [D] unexplained variation/residuals. All values [A] & [C] > 0 were significant (α = 0.05).

We confirmed our parsimonious models’ suitability for each genetic diversity index by testing all possible model combinations. The full model constructed using forward-selection was either the second-best (*H*_S_, Table 4) or best model (*H*_T_, Table 5; & *G*”_ST_, Table 6). For *H*_S_, the difference in AICC between the best and full models was essentially meaningless (delta = 0.20; Table 4).

**TABLE 4.**
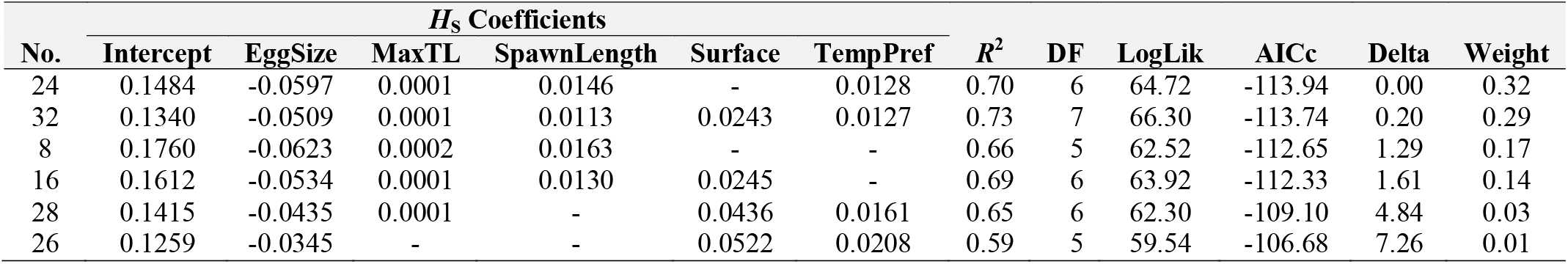
Model selection results for average site genetic diversity (*H*_S_) based on traits and diversity values measured for freshwater fish species (N=31; Table 1) collected across the White River Basin (Ozark Plateau, USA). Coefficients include intercept and a reduced set of traits (Table 2). No. = model number; *R*^2^ = coefficient of determination; DF = degrees of freedom; LogLik = log-likelihood; AICc = second-order Akaike Information Criterion; Delta = change in AICC from best model; Weight = model weight used for averaging.

| No. | $H_S$ Coefficients | | | | | | $R^2$ | DF | LogLik | AIC<sub>c</sub> | Delta | Weight |
| --- | --- | --- | --- | --- | --- | --- | --- | --- | --- | --- | --- | --- |
|  | Intercept | EggSize | MaxTL | SpawnLength | Surface | TempPref |  |  |  |  |  |  |
| 24 | 0.1484 | -0.0597 | 0.0001 | 0.0146 | - | 0.0128 | 0.70 | 6 | 64.72 | -113.94 | 0.00 | 0.32 |
| 32 | 0.1340 | -0.0509 | 0.0001 | 0.0113 | 0.0243 | 0.0127 | 0.73 | 7 | 66.30 | -113.74 | 0.20 | 0.29 |
| 8 | 0.1760 | -0.0623 | 0.0002 | 0.0163 | - | - | 0.66 | 5 | 62.52 | -112.65 | 1.29 | 0.17 |
| 16 | 0.1612 | -0.0534 | 0.0001 | 0.0130 | 0.0245 | - | 0.69 | 6 | 63.92 | -112.33 | 1.61 | 0.14 |
| 28 | 0.1415 | -0.0435 | 0.0001 | - | 0.0436 | 0.0161 | 0.65 | 6 | 62.30 | -109.10 | 4.84 | 0.03 |
| 26 | 0.1259 | -0.0345 | - | - | 0.0522 | 0.0208 | 0.59 | 5 | 59.54 | -106.68 | 7.26 | 0.01 |

**TABLE 5.**
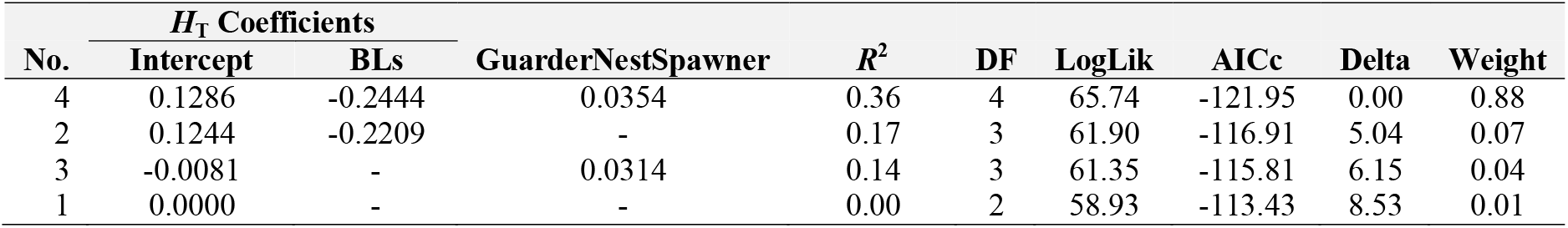
Model selection results for total genetic diversity (*H*_T_) based on traits and diversity values measured for freshwater fish species (N=31; Table 1) collected across the White River Basin (Ozark Plateau, USA). Coefficients include intercept and a reduced set of traits (Table 2). No. = model number; *R*^2^ = coefficient of determination; DF = degrees of freedom; LogLik = log-likelihood; AIC_c_ = second-order Akaike Information Criterion; Delta = change in AICC from best model; Weight = model weight used for averaging.

**TABLE 6.**
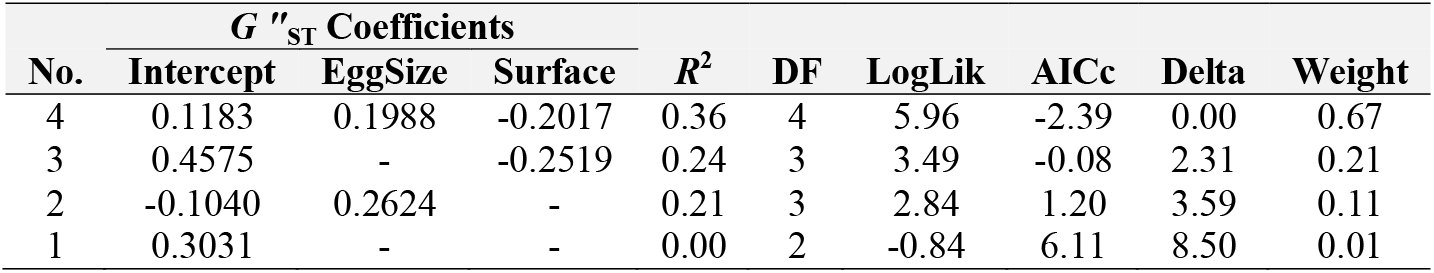
Model selection results for global genetic fixation/differentiation index (*G*”_ST_) based on traits and diversity values measured for freshwater fish species (N=31; Table 1) collected across the White River Basin (Ozark Plateau, USA). Coefficients include intercept and a reduced set of traits (Table 2). No. = model number; *R*^2^ = coefficient of determination; DF = degrees of freedom; LogLik = log-likelihood; AIC_c_ = second-order Akaike Information Criterion; Delta = change in AICC from best model; Weight = model weight used for averaging.

The averaged trait coefficients were used to predict genetic diversity indices for Slender Madtom. Predicted *H*_S_=0.10 with 95% confidence interval=0.07–0.13 (observed *H*_S_=0.16). Predicted *G*”_ST_=0.56 with 95% confidence interval=0.36–0.77 (observed *G*”_ST_=0.64). Unfortunately, *H*_T_ predictions are not comparable with observed values because the *residuals* of the linear relationship between *H*_T_ and N individuals were modeled rather than raw values.

## 4 DISCUSSION

Quantifying the relationship between traits and genetic diversity is one way to bridge the gap between ecology and evolution (Bohonak, 1999). Theoretically, traits that influence how genes are passed from one generation to the next, e.g., fecundity, or how genes are spread among populations, e.g., swimming ability, should impact genetic diversity (Papadopoulou & Knowles, 2016). Whether genetic diversity can be predicted from species traits alone is still an open question.

We found a significant signal of association between at least one genetic diversity index and half of the traits analyzed, thus supporting a hypothesized relationship between those traits and genetic diversity. Key morphological traits represented aspects of body size and swimming/foraging behavior: body elongation, lateral shape, pectoral fin position, caudal fin throttling, and position of the mouth. Important life-history traits included fecundity, egg size, length of spawning season, parental energy investment in offspring, and whether eggs were guarded post-spawn. The significant ecological traits were related to habitat use: Temperature preference, benthic feeding, and surface feeding.

### 4.1 Within-site genetic diversity (*H*_S_)

Species with higher within-site genetic diversity tended to be larger, with greater fecundity, smaller eggs, longer spawning seasons, preferences for warmer water temperature, with mouths phenotypically inclined toward the surface rather than the benthos (*vice-versa* for species with low diversity). The above relationships between genetic diversity and body size, egg size, environmental tolerance, and benthic *versus* pelagic spawning have been previously noted (Husemann et al., 2012; Osborne et al., 2014; Sousa-Santos et al., 2016).

Larger-bodied fish that feed more toward the surface and prefer warmer temperatures may occupy much larger areas than fish lacking such characteristics, i.e., dispersal syndromes (Comte & Olden, 2018). This may consequently increase both population sizes and connectivity, positively influencing genetic diversity within sites, and genetic diversity erodes more slowly in larger populations (Mitton & Lewis, 1989).

Population size and genetic diversity may also benefit from higher fecundity, smaller egg sizes, and extended spawning periods. Egg dispersal may also promote lotic genetic diversity through population connectivity, whereas those that produce more and smaller eggs may benefit from stream-mediated dispersal (Platania & Altenbach, 1998). The genetic diversity of fishes in lotic environments as it relates to egg dispersal is not well studied. However, it is consistent for marine fishes in that smaller eggs released into the water column disperse widely due to ocean currents (Riginos et al., 2014). Moreover, a more extended spawning season may provide greater environmental variability and, by chance or cue, provide optimal conditions for downstream dispersal and/or survival of eggs and larvae. For example, more turbulent flows during broadcast spawning may enhance viability by preventing egg clumping (Jager et al., 2001). Faster flows may also reduce competition by broadcasting embryos more widely and providing a more extensive range of ecological opportunities (McCabe Jr. & Tracy, 1994).

### 4.2 Total genetic diversity (*H*_T_)

Species with higher total genetic diversity tend to be deeper bodied, invest more energy into parental care, and construct/guard nests. This finding underscores the role of parental investment in determining overall genetic diversity. However, this finding stands in opposition to the suggestion that ‘*r*-strategists’ (high fecundity/low parental investment) tend to have larger population sizes and concomitantly higher genetic diversity than ‘*K*-strategists’ (low fecundity/high parental investment) (Mitton & Lewis, 1989; Romiguier et al., 2014). Although, those findings align more with our observation of within-site diversity above. Interestingly, a large meta-analysis of marine and freshwater fishes (*N*=463) also provided mixed support for the polarity of *r* vs. *K*-driven genetic diversity (Martinez et al., 2018). Romiguier et al. (2014) assessed 76 metazoan species across a diversity of evolutionary histories and observed an inverse relationship between propagule size (quality) and fecundity (quantity). Yet, we found fecundity positively correlated with higher parental investment and nest construction/guarding behavior. In our study species, those investing more in offspring also deposited more eggs - although smaller in size. Presumably, species exhibiting these characteristics also had greater numbers of offspring surviving to reproduce, thus promoting larger, more stable populations and higher genetic diversity. Ultimately, categorizing species as *r*- or *K*-strategists based on only a few traits may lead to spurious conclusions.

### 4.3 Among-site genetic diversity (*G*”_ST_)

Generally, species with higher among-site diversity tend to have mouths more ventrally positioned, trophically more benthic in orientation, and pectoral fins more dorsally positioned (for active swimming). Caudal fins were smaller (less influential in propulsion), whereas eggs were larger-sized (morphological interpretations from Brosse et al., 2021). Species with higher among-site diversity also seemingly displayed reduced within-site diversity. Indeed, these characteristics seem opposed to those promoting higher within-site diversity (as above). Species in the latter category are more benthic-oriented (per mouth and pectoral fin positions), with trophic resources gleaned from the bottoms of streams/rivers. Hence, they display less movement than heterospecifics trophically oriented within the water column or the surface. Moreover, their larger eggs are perhaps less likely to be dispersed downstream. These results are consistent with the expectation that benthic habitat specialists demonstrate greater divergences among populations than habitat generalists (Pilger et al., 2017).

### 4.4 Can genetic diversity be predicted?

The processes that have shaped global biodiversity must be clearly understood before attempting to mitigate its loss (Manel et al., 2020). In this regard, threatened taxa lose heterozygosity more rapidly due to genetic drift acting on declining populations. Concomitantly, their genetic diversities are depressed (Spielman et al., 2004), which imparts a substantial, adverse effect on fitness (DeWoody et al., 2021). If managers could identify which species tend to have lower levels of genetic diversity, they could prioritize them for targeted surveys and subsequent management. The capacity to forecast which species have lower genetic diversity based on easily estimated traits would provide much-needed focus, as there are far too many species for each to be evaluated for population genetic metrics.

While a significant relationship between traits and genetic diversity is of interest, it may be of scant applicability if traits lack predictive power. For example, deterministic trait-based models are of little help if genetic diversity variation among species is stochastically-driven. However, previous studies suggest that a trait-based framework may be valuable. Meirmans et al. (2011) found that ecological and life-history traits explained 30% of the variation in the genetic structure of alpine plant species, a remarkably high value given it was based on but six characters, with other (non-assayed) processes also influential. Other studies across a diverse group of animals identified even higher correlations (e.g., *R*^2^=0.79) between genetic diversity and either life-history traits (Romiguier et al., 2014) or dispersal abilities (Bohonak, 1999). By comparison, our models of genetic diversity performed well with adj. *R*^2^ ranging from 0.31 to 0.68. We acknowledge potential drawbacks in evaluating the predictive capacity of models solely based on coefficients of determination (Onyutha, 2020). In particular, our within-population genetic diversity model (*H*_S_) provided the best predictive capacity (Std Err=0.016), whereas our among-population genetic diversity model (*G*”_ST_) had a greater standard error (Std Err=0.11). However, both reasonably predicted *H*_S_ and *G*”_ST_ for the Slender Madtom. More validation is needed to determine how useful these models can indeed be.

### 4.5 Phylogenetic autocorrelation

Genetic diversity is not a heritable trait of a species *per se* but rather an emergent property (Duminil et al., 2007). We thus hypothesize that similarity in genetic diversity among related species is driven by species-specific traits that mediate vital components of genetic diversity, such as population sizes and connectivity (Abrams, 2019; Fobert et al., 2019; Naish et al., 2013). Since closely related species tend to be more similar with respect to traits (Felsenstein, 1985), the same might be expected for genetic diversity in that it emerges via trait combinations (Duminil et al., 2007). We thus suggest that species with similar traits should reflect similar genetic diversities, although the mechanism is not directly due to shared history (independent of traits).

To avoid overestimating significance due to autocorrelation, we considered the effects of shared evolutionary history in testing the overall relationship between genetic diversity and traits (Hawkins, 2012). However, we did not incorporate phylogeny in predictive modeling (Meirmans et al., 2011). Strictly speaking, our study species are not independent of one another, and our statistical application of linear models would violate the assumption of independence. However, we would also remove favorable aspects of statistical signal by forcing independence by factoring out variance due to shared ancestry (Duminil et al., 2007). We agree with others (Legendre, 1993; Peres-Neto, 2006) that a balanced approach to autocorrelation is necessary. In this sense, phylogenetic autocorrelation is interpreted not as bias or *artifact* but instead as what we are interested in (Hawkins, 2012; Legendre, 1993).

## 5 Conclusion

Although all estimators (i.e., *H*_T_, *H*_S_, and *G*”_ST_) employed herein are mathematically related, we found that each captured different facets suggesting mechanisms by which life historic and other phenotypic characters relate to genetic diversity. Traits evolve over deep and shallow timeframes as environments differentiate and populations interact and diverge. Although viewed as conduits between ecology and evolutionary history, the association of these traits has been difficult to establish unequivocally. However, if so verified, the link between ecology and evolution can be estimated by modeling appropriate traits. Genetic diversities could then be quantified for species deemed ‘sibling’ (i.e., near-identical morphologically), ‘cryptic’ (as previous but non-hybridizing), or with narrow niches/restricted distributions (e.g., short-range endemics; Davis et al., 2015). Thus, forecasting genetic diversity would be valuable for deriving conservation policy and subsequently applying management decisions (Hoban et al., 2022). An approach is needed to focus and prioritize management and conservation, particularly given the multiplicity of species spanning numerous distinct geographic regions. The procedure described herein can facilitate management and streamline conservation plans by deriving diversity metrics for freshwater species currently lacking such information. Expanding the trait-based approach by incorporating more species, regions, greater phylogenetic breadth, and numerous traits will undoubtedly provide more focused insights and possibly greater predictive power. Furthermore, this community genomics approach should incorporate biotic interactions among species (*sensu* Hand et al., 2015) to integrate an overlooked essential biotic component. The framework herein also provides a platform for bridging the gap between micro- and macro-evolution, in that traits impinging upon genetic diversity (microevolution) could also play a significant role in speciation and extinction (macroevolution) (Singhal et al., 2018).

## ACKNOWLEDGEMENTS

We are grateful to M. Flurry, M. George, T. Goodhart, K. Hollar, and M. Reed for assisting with DNA extractions. The Arkansas High-Performance Computing Center provided analytical resources. Funding was provided by the University of Arkansas Distinguished Doctoral Fellowship (TKC and ZDZ), the Harry and Jo Leggett Chancellor’s Fellowship (ZDZ), the Bruker Professorship in Life Sciences (MRD), the Twenty-First Century Chair in Global Change Biology (MED), and an NSF Postdoctoral Research Fellowship in Biology (TKC) [DBI: 2010774]. The findings, conclusions, and opinions expressed in this article represent those of the authors and do not necessarily represent the views of the NSF nor other affiliated or contributing organizations.

## AUTHOR CONTRIBUTIONS

ZDZ conceived the research with input from all authors. Specimen collection was done by ZDZ & TKC. ZDZ did laboratory work, bioinformatics, data analysis, and manuscript drafting. All authors contributed to the interpretation of results, formulating conclusions, and critically revising the manuscript. MRD and MED administered funding through their University of Arkansas Endowments.

