## Supplemental Material for "Trait-based predictors of genomic diversity within freshwater fish communities of the Ozarks"

**Traits and relatedness associated with variability of genetic diversity among freshwater fish species of the Ozarks**

**TABLE OF CONTENTS**

| Supplement S1 | Fish Species Trait Info | p. 1 |
| --- | --- | --- |
| Supplement S2 | Histograms/distributions of genetic variables and covariates | p. 2 |
| Supplement S3 | Histrograms/distributions of trait variables | p. 3 |
| Supplement S4 | PCA of fish species genetic variables | p. 4 |
| Supplement S5 | Plot of Hs and correlated traits | p. 5 |
| Supplement S6 | Plot of Ht and correlated triats | p. 6 |
| Supplement S7 | Plot of Gst and correlated traits | p. 7 |

**SUPPLEMENT S1.** Fish species collected across the White River Basin of the Ozarks, USA (N=31) were defined by characteristics (N=28) broadly related to morphology, life history, and ecology. These traits were gathered from a vareity of sources shown here.

| **Code** | **Trait** | **Units** | **Source** | **Notes** |
| --- | --- | --- | --- | --- |
| MaxTL | Max body length | millimeters | Olden; FishMorph; FishTraits; FoA | Deferred to Olden |
| LengMature | Mean length at maturity | millimeters | Olden |  |
| AspectRatio | Caudal fin aspect ratio | height^2^/surface area | Olden |  |
| BEl | Body elongation | length/depth | FishMorph |  |
| VEp | Vertical eye position | eye position/body depth | FishMorph |  |
| REs | Relative eye size | eye diameter/head depth | FishMorph |  |
| OGp | Oral gape position | mouth height/body depth | FishMorph |  |
| RMl | Relative maxillary length | maxillary length/head depth | FishMorph |  |
| BLs | Body lateral shape | head depth/body depth | FishMorph |  |
| PFv | Pectoral fin vertical position | pectoral fin vert position/body depth | FishMorph |  |
| PFs | Pectoral fin size | pectoral fin length/body length | FishMorph |  |
| CPt | Caudal peduncle throttling | caudal fin depth/caudal peduncle depth | FishMorph |  |
| AgeMature | Age at maturation | years | Olden; FishTraits; FoA | Deferred to Olden |
| Longevity | Longevity | years | Olden; FishTraits; FoA | Deferred to Olden |
| Fecundity | Fecundity | number eggs/season/female | Olden; FishTraits; FoA | Deferred to Olden |
| EggSize | Mean egg diameter | mm | Olden |  |
| SpawnLength | Length of spawning season | months | FishTraits |  |
| NonGuardBroodHide | Non-guarding brood hiders | binary | Olden; FishTraits; FoA |  |
| NonGuardOpenSubstrate | Non-guarding substrate spawners | binary | Olden; FishTraits; FoA |  |
| GuarderNestSpawner | Egg Guarding nest spawners | binary | Olden; FishTraits; FoA |  |
| SpawnFreq | Spawning frequency | binary | Olden; FishTraits |  |
| ParentEnergy | Parental Care Index | discrete index | Olden |  |
| DietIndex | Diet Generality Index | discrete index | Derived from FishTraits | Sum of binary diet variables |
| HabitatIndex | Habitat Generality Index | discrete index | Derived from FishTraits | Sum of binary habitat variables |
| TrophGuild | Trophic guild | categorical | Olden |  |
| TempPref | Water temperature preference | categorical | Olden |  |
| Benthic | Benthic dwelling | binary | Olden; FishTraits; FoA | Deferred to Olden |
| Surface | Surface dwelling | binary | FishTraits |  |
| **Olden =** |  |  |  |  |
| Julian D. Olden (2022, unpublished data) | | | | |
| Mims, M.C., Olden, J.D., Shattuck, Z.R., and N.L. Poff (2010). Life history trait diversity of native freshwater fishes in North America. *Ecology of Freshwater Fish* 19:390-400 | | | | |
| Giam X, Olden JD. (2016). Environment and predation govern fish community assembly in temperate streams. *Global Ecology and Biogeography* 25: 1194-1205 | | | | |
| **FishMorph =** |  |  |  |  |
| Brosse, S., Charpin, N., Su, G., Toussaint, A., Herrera‐R, G. A., Tedesco, P. A., & Villéger, S. (2021). FISHMORPH: A global database on morphological traits of freshwater fishes. *Global Ecology and Biogeography* 30(12), 2330-2336. | | | | |
| **FishTraits =** |  |  |  |  |
| Frimpong, E. A., & Angermeier, P. L. (2009). Fish traits: a database of ecological and life-history traits of freshwater fishes of the United States. *Fisheries* *34*(10), 487-495. | | | | |
| **FoA =** |  |  |  |  |
| Robison, H. W., & Buchanan, T. M. (2020). *Fishes of Arkansas*. University of Arkansas Press. | | | | |

**SUPPLEMENT S2.** Three statistics summarizing genetic diversity (HS, HT, and G”ST) and four covariates were calculated for fish species (N=31) collected across the White River Basin of the Ozarks, USA. Nindiv = the total number of individuals genotyped/analyzed; Nsites = total number of sites where genotyped/analyzed individuals were collected; Median_RivDist = the median value of pairwise river network distance between sites at which each species occurred; Mean_Ind_Site = mean number of individuals genotyped/analyzed per site.

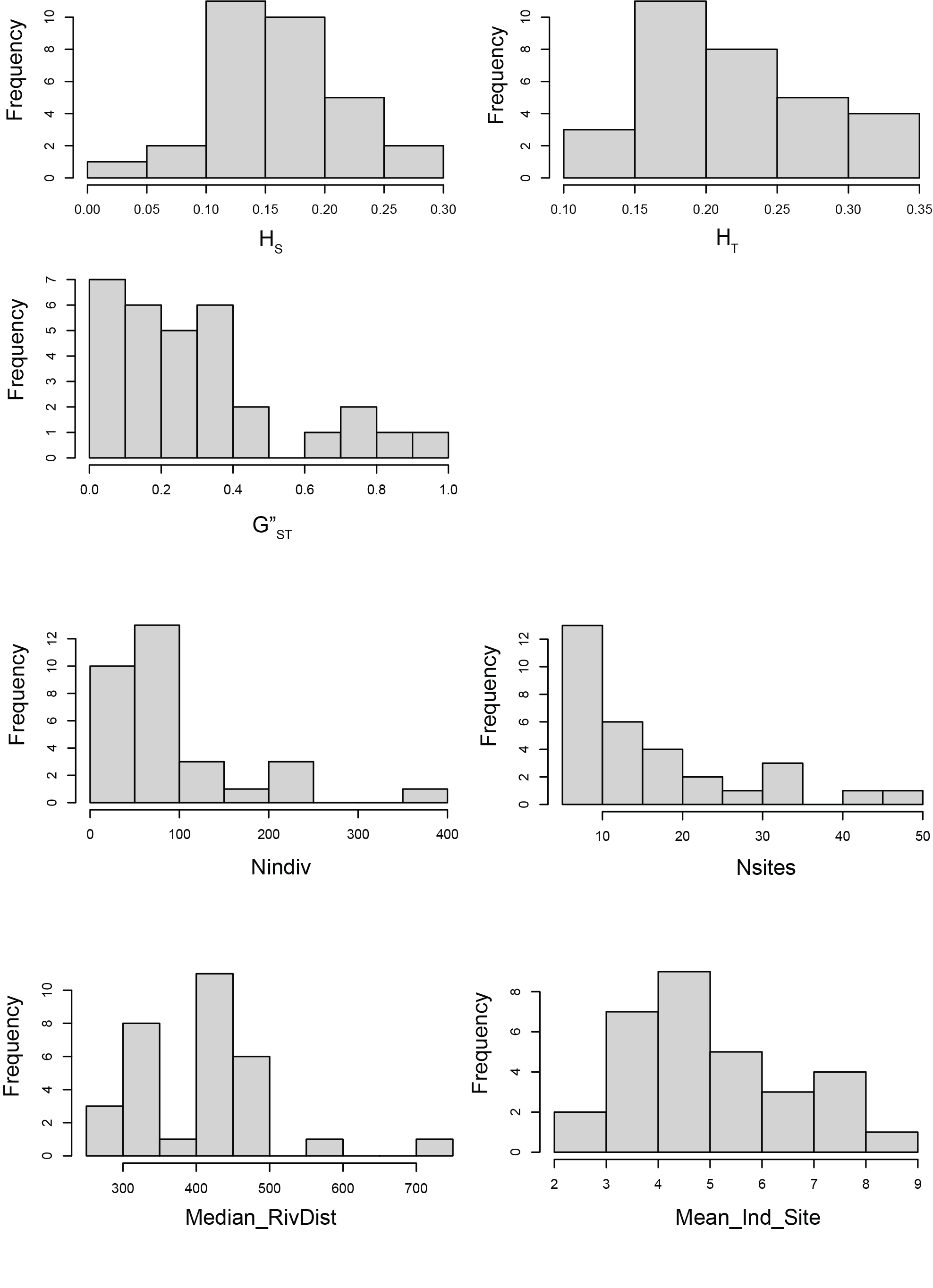

**SUPPLEMENT S3.** Principal components analysis of three genetic summary variables quantified for thirty-one fish species collected across the White River Basin of the Ozarks, USA. *H*_S_ = heterozygosity within sites; *H*_T_= total heterozygosity; *G”*_ST_ = genetic differentiation/fixation coefficient.

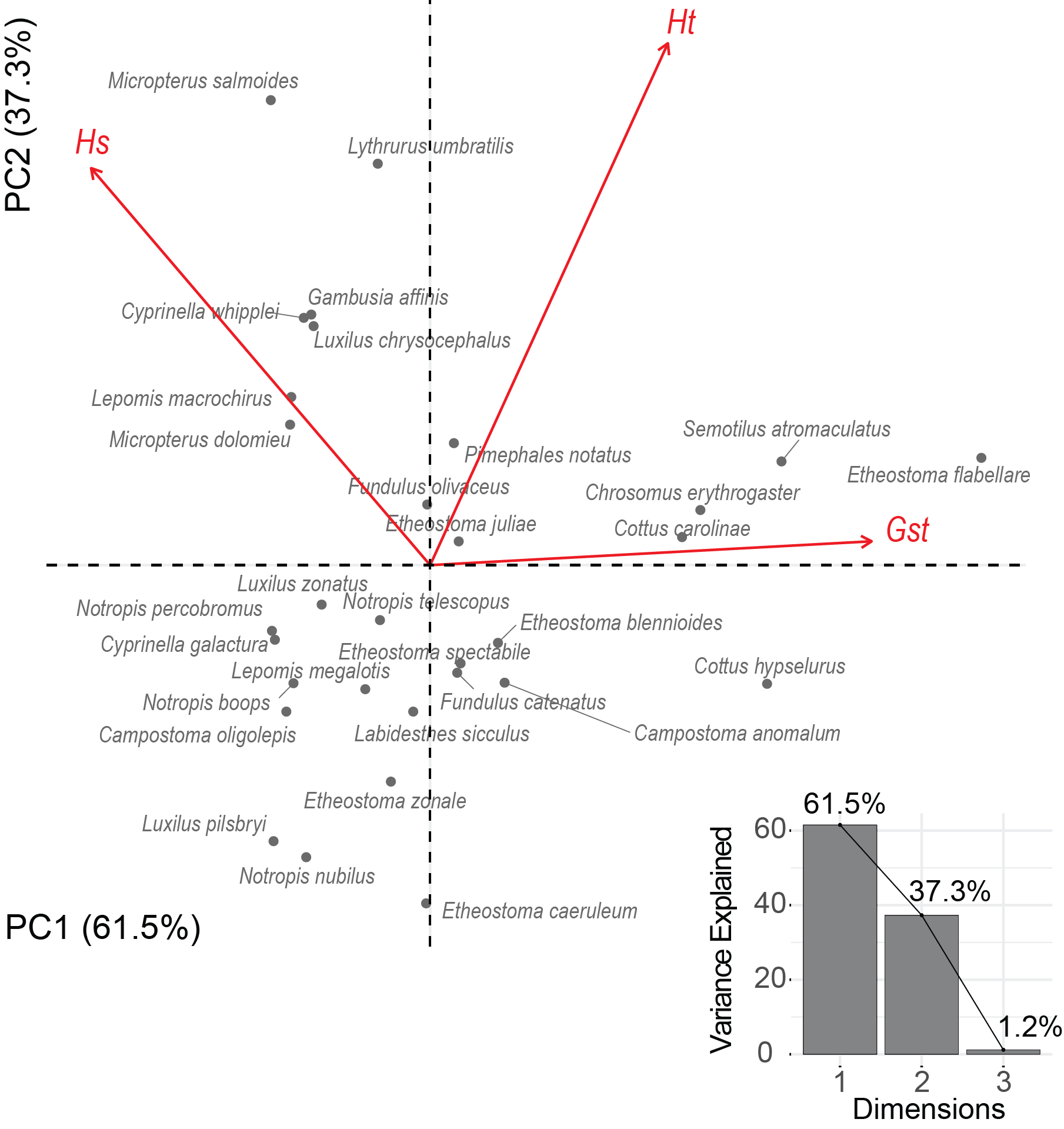

**SUPPLEMENT S4.** These histograms show distributions of twenty-eight trait measurments for thrity-one fish species collected across the White River Basin of the Ozarks, USA. Traits were not transformed so as to maximize interpretability of predictive models.

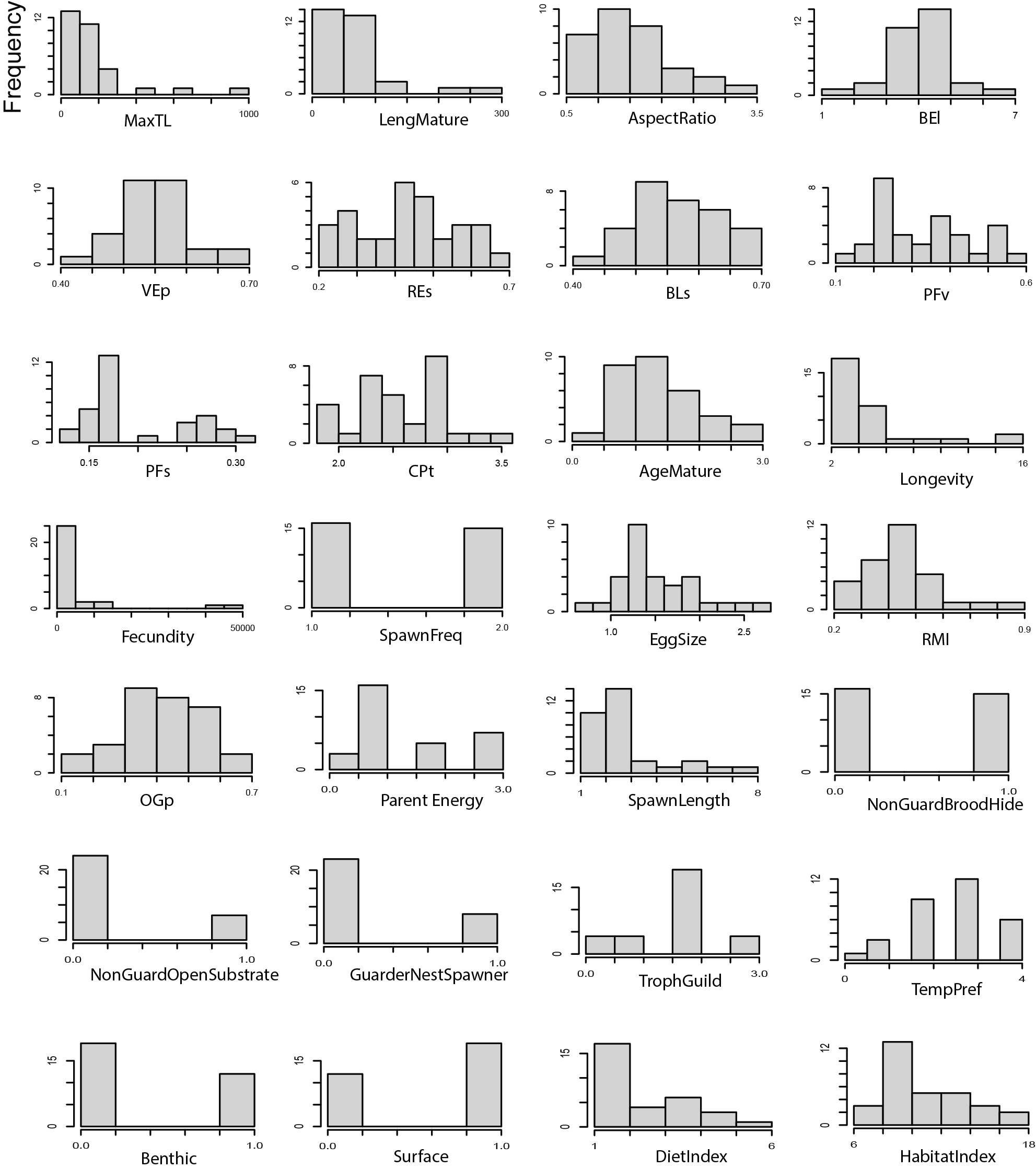

**SUPPLEMENT S5.** Traits showing significant correlation with average site genetic diversity (*H*_S_) quantified for *N=*31 freshwater fish species collected across the White River Basin of the Ozarks, USA.

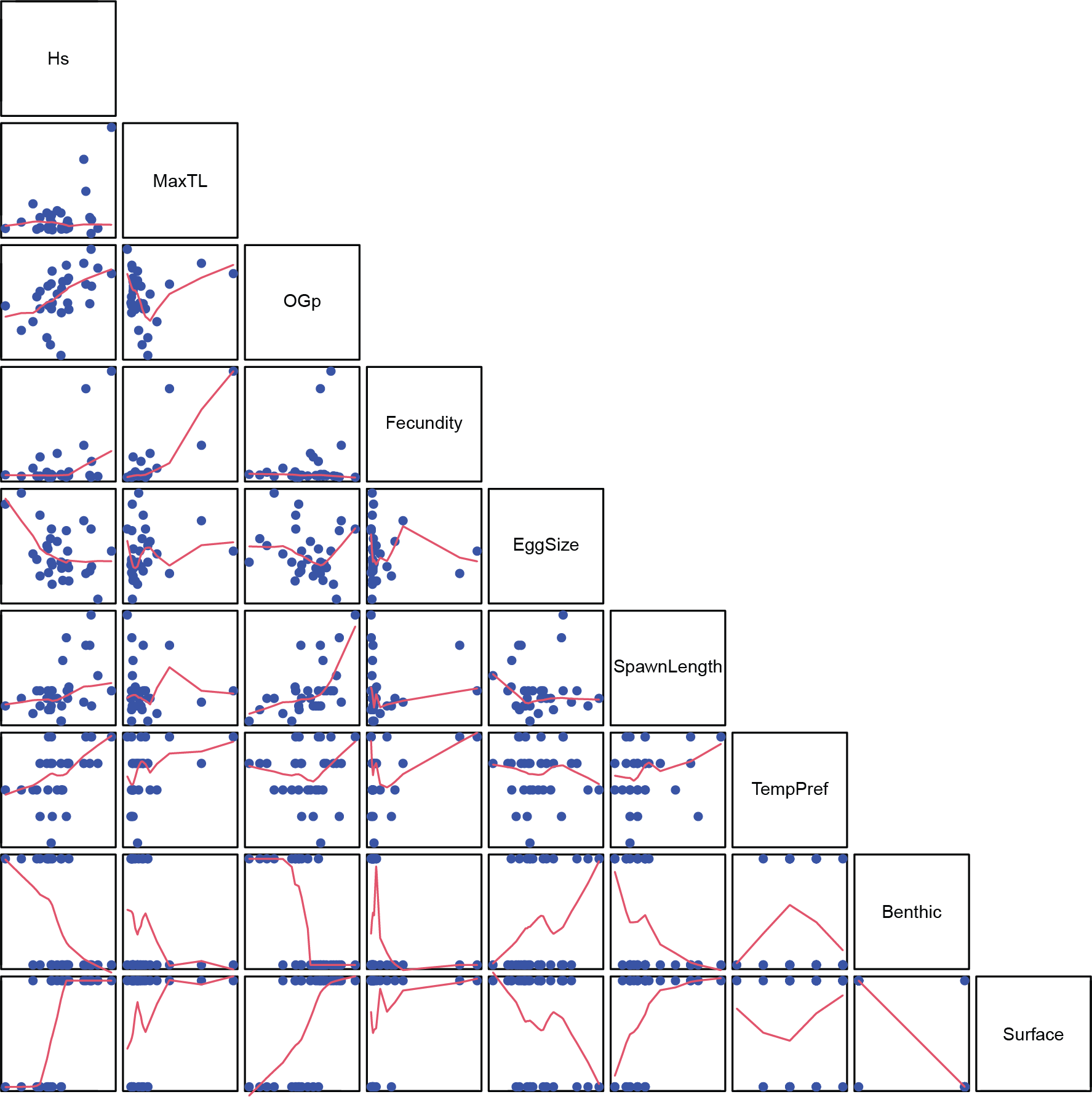

**SUPPLEMENT S6.** Covariates and traits showing significant correlation with total genetic diversity (*H*_T_) quantified for *N=*31 freshwater fish species collected across the White River Basin of the Ozarks, USA. Note: traits were related to the residuals of the relationship between *H*_T_ the covariates.

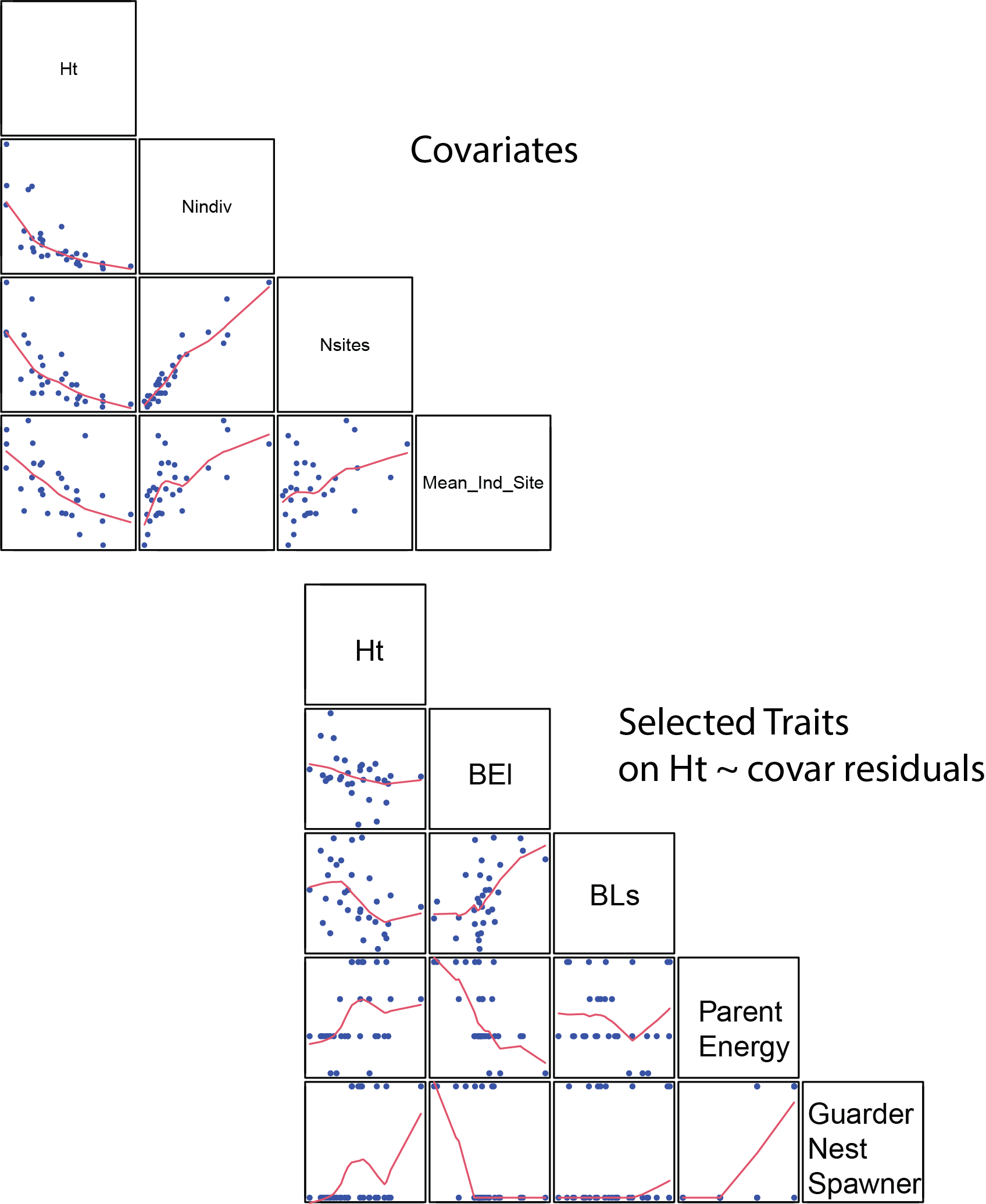

**SUPPLEMENT S7.** Traits showing significant correlation with global genetic fixation/differentiation (*G”*_ST_) quantified for *N=*31 freshwater fish species collected across the White River Basin of the Ozarks, USA.

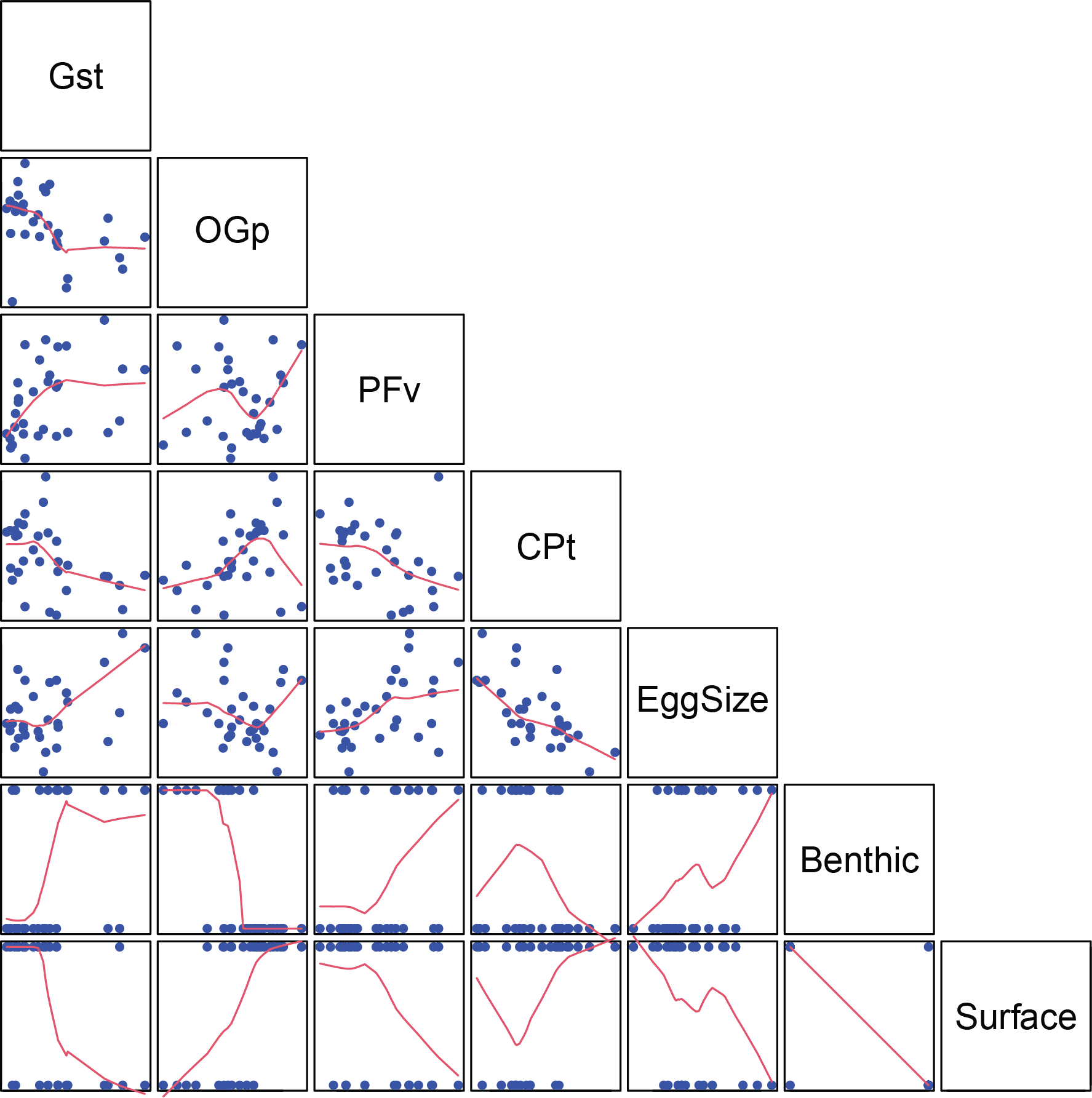
